# Scrounging by foragers can resolve the paradox of enrichment

**DOI:** 10.1101/077537

**Authors:** Wataru Toyokawa

## Abstract

Theoretical models of predator-prey system predict that sufficient enrichment of prey can generate large amplitude limit cycles, paradoxically causing a high risk of extinction (the paradox of enrichment). While real ecological communities contain many gregarious species whose foraging behaviour should be influenced by socially transmitted information, few theoretical studies have examined the possibility that social foraging might be a resolution of the paradox. I considered a predator population in which individuals play the producer-scrounger foraging game both in a one-prey-one-predator system and a two-prey-one-predator system. I analysed the stability of a coexisting equilibrium point in the former one-prey system and that of non-equilibrium dynamics of the latter two-prey system. The result showed that social foraging can stabilise both systems and thereby resolves the paradox of enrichment when scrounging behaviour is prevalent in predators. This suggests a previously neglected mechanism underlying a powerful effect of group-living animals on sustainability of ecological communities.

## 1 Introduction

Understanding how complex biological communities can sustainably persist has been an essential theme in ecology. Community ecologists have tried for decades to reveal mechanisms that maintain or destroy the persistence of natural communities [1–3]. One of the most intriguing predictions from classical predator-prey models is that a sufficient enrichment of prey (i.e., increasing prey carrying capacity) can destabilise the community, causing a high risk of extinction [2]. This is called the paradox of enrichment. Although this hypothesis was supported in simple predator-prey systems [4–6], it has been rejected by a number of empirical studies (e.g. [7–10]).

The bulk of theoretical studies have identified ecologically relevant mechanisms that can explain the scarcity of the paradox of enrichment in field conditions. Generally speaking, mechanisms that reduce the per predator consumption rate as predator density increases are thought to weaken the paradox of enrichment [11]. For instance, both inducible defensive morphs or predator-avoidance behaviour in prey [12,13], and aggressive mutual interference or “prudence” in predators [14, 15] are depicted as potential resolutions of the paradox in simple predator-prey systems. Also, as for more complex communities including multiple prey populations, theory predicts that imperfection of the optimal menu switching by predator [16] and diversity in interaction types between species [17] can help the community to persist.

However, most of the previous predator-prey studies have considered an asocial forager that searches for food resources without using socially transmitted information. Assuming asocial foraging is simple and mathematically tractable, and it might be a suitable assumption for some communities such as phytoplankton-zooplankton systems (e.g. [6]). However, ecological communities actually contain many gregarious species, whose foraging behaviour should be influenced by information that comes from other conspecifics.

Scrounging (or kleptoparasitism) is a well-known consequence of social information use in predation [18]. Assume that an individual in a group engages in predation by using either of two different tactics: searching their environment for food clumps (or divisible prey) by itself (”producing”), or attending to other foragers’ clump discoveries and sequestering some food at each clump found by a producer (”scrounging”). Note that scrounging does not require any aggressive interferences between predators, and can occur even in mere aggregations of animals which do not consist of social structures or genetic relations between individuals [18]. Rather, social foraging can potentially exist in any circumstances under which animals search for divisible food and information that an individual has found/captured a food clump is available to other conspecifics. For this reason, social foraging should be a very common phenomenon in ecological communities.

The game theoretic model of this producer-scrounger (PS) behavioural dynamics predicts that both producer and scrounger tactics can stably coexist at the equilibrium [18–22]. The equilibrium proportion of the two tactics may have a substantial influence on the predator-prey dynamics, because prey are discovered only by producing predators. The proportion of producers in the predator population should, therefore, crucially affect the predation pressure (i.e., predator-prey encounter rate).

Conversely, population dynamics may affect the equilibrium proportion between producers and scroungers. In the basic PS game, each producer exclusively obtains a “finder’s advantage”, that is a portion *f* out of the total value of a prey item *F*(0 ≤ f ≤ *F*), before scroungers arrive. Once a producer capture a prey, all of the scroungers arrive and divide the remaining *F − f* value equally among the individuals present (i.e., all the scroungers and the producer). Scrounging provides more rewards when scroungers are rare (i.e., producers are common) while it becomes less rewardable when they are common. This negative frequency dependence generates an evolutionary stable mixed equilibrium [23] at which the producers’ proportion is *q*∗ = *f/F + 1/G*, where *G* is a group size [19]. Note that the equilibrium proportion of producers is reduced by an increase in the group size *G* which might be influenced by the population size. Therefore, the model predicts that the proportion of producing would be influenced not only by the finder’s advantage, but also by population dynamics.

Although both the paradox of enrichment and producer-scrounger dynamics have separately received huge attention by ecologists, the relationship between them remains largely unclear. A notable exception was a Coolen et al. [24] study, which showed that scrounging behaviour in predators can stabilise the oscillation of the classical Lotka-Volterra predator-prey model. As mentioned above, an increase of predator population density (i.e., group size) reduces the proportion of producing, and reduction of producer proportion should reduce the per capita predation rate. Therefore, predation pressure becomes mitigated as the predator population grows, and thereby the Lotka-Volterra system is stabilised [24]. Although the Lotka-Volterra model is too simple to understand complex communities and does not contain the paradox of enrichment because of the absence of prey carrying capacity, Coolen et al. [24] suggests that the PS game in predator is a strong candidate for the resolution of the paradox of enrichment in more complex predator-prey systems.

In this article, I extend the model of Coolen et al. [24] to standard one-prey-one-predator systems (i.e., Rosenzweig-MacArthur model [1]) as well as more complex two-prey-one-predator systems [25–28]. Both of these systems are known to show the paradox of enrichment. Here I demonstrate that the producer-scrounger dynamics in the predator resolves the paradox under a broad range of conditions.

## 2 One prey - one predator system

### 2.1 The model

First, I investigated a standard predator-prey model consisting of a prey population *X* and a predator population *Y* [1]. I followed the assumptions in Coolen et al. [24] to model PS game dynamics in predators as follows. I supposed that the predator population is divided into *g* (*g* ∈ {1, 2, 3,…}) groups of G individuals each (*Y* = *gG*), in which individuals play the PS game with other conspecifics. Predator individuals can search for food using either producing (i.e., searching for food asocially and capturing it by itself) or scrounging (i.e., wait for other individuals to find food and then kleptoparasitize it) ^1^. Each producer obtains a finder’s advantage *f* out of *F* energetic units (0 *≤ f ≤ F*), before the scroungers arrive. Once a producer captures a prey item, all of the scroungers arrive and divide the remaining *F − f* energetic units equally among the individuals present. The proportion of producers among the predators is *q* (0 *≤ q ≤* 1). I assumed that prey discovery is rare enough to occur sequentially, so that all scroungers can visit all events of prey discoveries by producers. Under these assumptions, the evolutionary or behaviourally stable equilibrium of producer proportion is *q*\* = *g/Y* + *f/F* [18,19]. Note that, because the proportion *q*\* should not be larger than 1, the model is ecologically relevant when it satisfies 1 ≤ *g ≤ Y* (1 *− f/F*). I further assumed that behavioural plasticity (e.g., learning) allow individuals to adjust to the PS game’s equilibrium *q\** within a time scale much shorter than predator birth and death processes [29], so that *q* is always equals to *q\** in the following one-prey-one-predator dynamics:

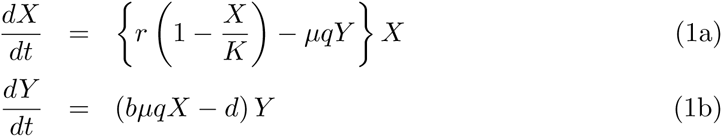

where *µ* = *a*/(1 + *ahX*) and *q* = *q*\* = *g/Y + f/F*.

*µ* implies the instantaneous per capita capturing rate of prey, depicted by a Holling type II functional response with searching efficiency *a* and handling time *h*; *b* is the conversion efficiency, which relates the predator’s birth rate to prey consumption, and *d* is the death rate of the predator. Note that producers (*q*Y*), but not scroungers, are influencing the number of prey captured. For prey population *X*, *r* is the per capita intrinsic growth rate, and *K* is the carrying capacity which traditionally indicates the degree of enrichment [2,30,31].

### 2.2 Method

To analyse the effect of scrounging behaviour in predators on the paradox of enrichment, I focused on the proportion of a finder’s advantage *f/F* [24]. As seen in the equation *q*\* = *g/Y + f/F*, *f/F* determines the equilibrium proportion of producers, and hence that of scroungers. The smaller the finder’s advantage is, the more prominent the effect of scrounging should be.

I analysed the local stability around a coexisting equilibrium point at which both prey and predator densities are positive. It is well known that the local stability can be analysed graphically in the predator-prey phase plane: In the classical Rosenzweig-MacAurthur model, the equilibrium is stable if the vertical predator isocline (*dY/dt* = 0) intersects with the right-hand side of the humped prey isocline (*dX/dt* = 0), while it becomes unstable when predator isocline intersects the left of the hump [1,32]. Because an increase in *K* does not affect the predator isocline, the increase in *K* will eventually place the predator isocline left of the hump, making the equilibrium unstable (the paradox of enrichment). Here, I present the similar graphical analysis to show the relation between the equilibrium stability and prey enrichment.

### 2.3 Result

Figure 1A and 1B show both predator and prey isoclines of the model (Eq. 1). When the finder’s advantage is sufficiently small (i.e, *f/F ≤ dh/b*), the predator isocline is concave-down and never intersects with the left side of the humped prey isocline (Figure 1A, Appendix A), and thereby the predator and prey can stably coexist regardless of prey enrichment. Figure 1A also shows that the equilibrium densities of both species increase with enrichment when *f/F ≤ dh/b*. Therefore, the paradox of enrichment disappears if scrounging behaviour is prominent in the predator population.

**Figure 1.**
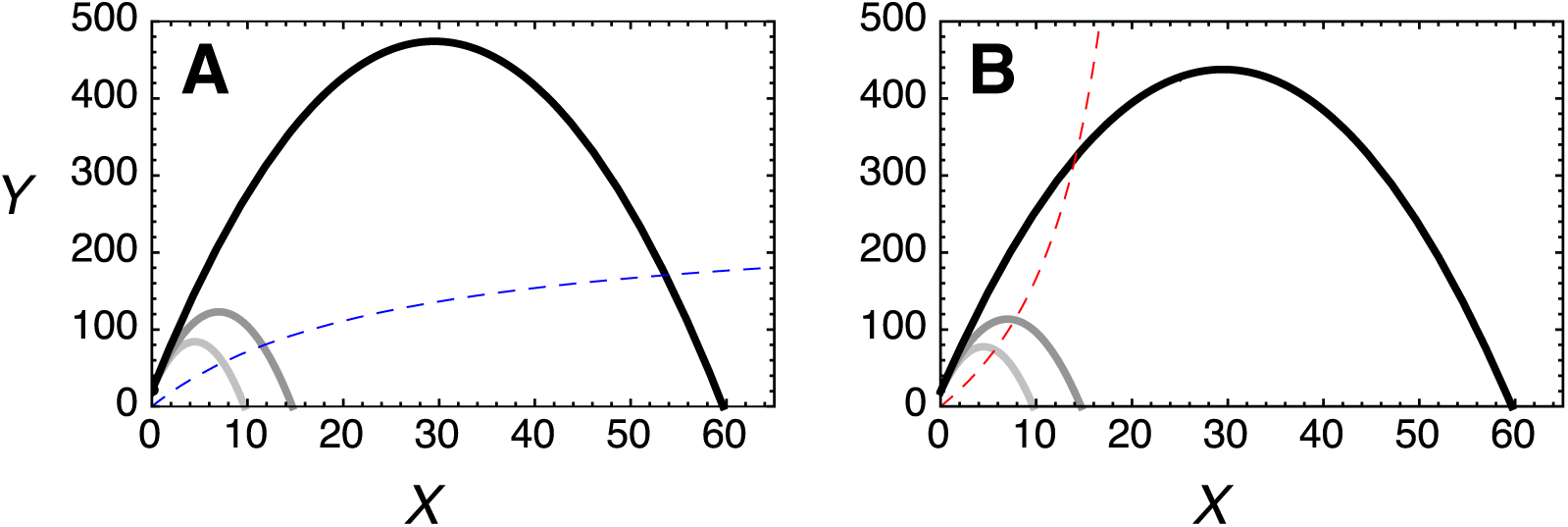
Phase-plane diagrams. Hump shaped solid lines are prey isoclines with different carrying capacities (light grey: *K* = 10, grey: *K* = 15, black: *K* = 60). Dashed lines are predator isoclines, when (A) the finder’s advantage is small (*f/F ≤ dh/b*) and (B) the finder’s advantage is large (*f/F > dh/b*). The intersects of the both isoclines are coexistence equilibria. Predator isocline is the same for all carrying capacity levels. Parameters were set to the following values: *r* = 15, *a* = 1, *b* = 0.5, *h* = 1, *d* = 0.25, *F* = 1, *g* = 5, *f* = 0.48 for A and *f* = 0.52 for B.

On the other hand, when the finder’s advantage is large (i.e, *f/F > dh/b*), the predator isocline is concave-up and the intersect eventually move to the left side of the hump as *K* increases, and hence the paradox still exists (Figure 1B).

## 3 Two prey - one predator system

### 3.1 The model

My next model is a familiar two-prey-one-predator system in which the predator practices optimal foraging based on the profitabilities and abundances of two prey (e.g. [16, 25–28]). Genkai-Kato and Yamamura [26] studied non-equilibrium dynamics of the basal model, and found that the profitability of the less-profitable prey regulates the amplitude of population oscillations. Nevertheless, the system is globally unstable and the paradox of enrichment remains prominent under a range of conditions. Assuming the same PS game in the predator population of the basal model, I investigated the effect of scrounging on the non-equilibrium dynamics of this system.

This two-prey-one-predator system, which consists of more-profitable prey *X*_1_, less-profitable prey *X*_2_, and predator *Y*, is described as follows:

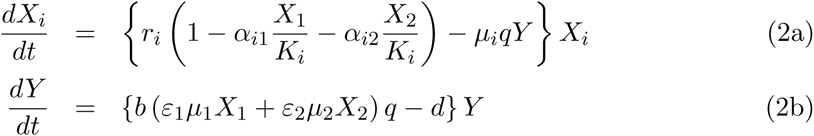

where 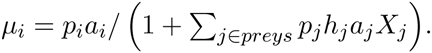

For predator population *Y*, *µ*_i_ implies the instantaneous per capita capture rate for prey *i* (*i* ∈ {1, 2}), depicted by a Holling type II functional response; *q* (0 *≤ q ≤* 1) is a proportion of producers in the predator population; *a*_*i*_ is a searching efficiency for prey *i*; *h*_*i*_ is the handling time for prey *i*; *ε*_*i*_ is the energy value of an individual of prey *i*; and *p*_*i*_ (0 *≤ p_i_ ≤* 1) is the capture probability of an individual of prey *i* given an encounter; *b* is the conversion efficiency, which relates the predator’s birth rate to prey consumption, and *d* is the death rate of the predator species. For prey *i*, *α*_*ij*_ are the intra- and interspecific competition coefficients (*α*_*ii*_ = 1); *r*_*i*_ is the per capita intrinsic growth rate of prey *i*; and *K*_*i*_ is the carrying capacity of prey *i*.

Assume that the predator is an optimal forager that chooses the value for each of the probabilities *p*_*i*_ in order to maximise the energy input by predation *ε*_1_*µ*_1_*X*_1_ + *ε*_2_*µ*_2_*X*_2_. I also assume that the prey *X*_1_ is more profitable for the predator than prey *X*_2_, i.e., *ε*_1_/*h*_1_ > *ε*_2_/*h*_2_ so that *p*_1_ should always be 1 (i.e., prey 1 is always included in the diet [33]), while the more-profitable prey *X*_1_ is superior in competition to the less-profitable prey (*α*_12_ < *α*_21_).

#### 3.1.1 Optimal foraging

I assumed that the predators are optimal foragers that select their diet in accordance with optimal diet utilisation theory [34]. Under perfect optimal foraging, the capture probability of an individual of the less-profitable prey given an encounter, *p*_2_, equals zero or one, depending on whether the density of the more-profitable prey is greater or less than the threshold density 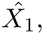, where 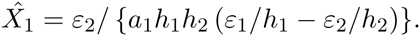. If the density of the more-profitable prey drops below this critical threshold (i.e., the diet-change threshold), the less-profitable prey is also included in the diet (*p*_2_ = 1). Otherwise, the less-profitable prey is excluded from the diet (*p*_2_ = 0). Inclusion or exclusion of the less-profitable prey depends on the difference in profitability (i.e., the expected energy values per handling time *ε/h*) and the density of the more-profitable prey [34].

#### 3.1.2 Producer-scrounger game under multiple prey types

Because there are two different prey populations, I had to consider the PS game under multiple food types. As in the one-prey-one-predator model described above, I suppose that the predator population (*Y*) is divided into *g* groups of *G* individuals each (*Y* = *gG*), and the predator individuals can choose either producing or scrounging. In my two-prey-one-predator system (Eq. 2), the instantaneous total number of prey captured by a single producer is *µ*_1_*X*_1_ + *µ*_2_*X*_2_. As in the classic PS game model [19], a producer individual capturing a prey *i* obtain a finder’s advantage *f*_*i*_ out of the maximum *F*_*i*_ energy available before the arrival of scroungers, and then the remaining *F*_*i*_−*f*_*i*_ energetic units are equally divided between the producer and all scroungers present in the group. I also assumed again that all scroungers can visit all events of prey discoveries by producers in their group. Expected instantaneous per capita energy intake of both producer (*I*_*p*_) and scrounger (*I*_*s*_) are given by:

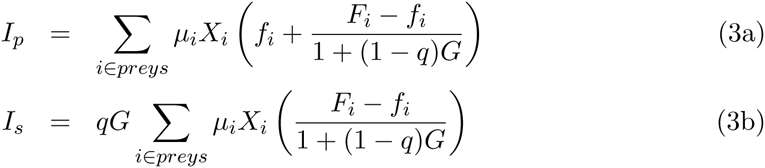

where *q* (0 ≤ *q* ≤ 1) is the proportion of producers.

Setting *I*_*p*_ = *I*_*s*_ obtains a behaviourally (or evolutionally) stable strategy [23] of producing probability *q\** (Appendix B):

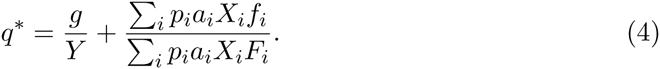

Note that when *F* = *F*_1_ = *F*_2_ and *f* = *f*_1_ = *f*_2_, the equilibrium is consistent with the original PS game equilibrium *q\** = *g/Y* + *f/F* [18]. Following the one-prey-one-predator model, I further assume that behavioural plasticity allows predator individuals to adjust to the behavioural ESS within the single time step of the population dynamics, so that they always achieve *q* = *q\** at the timescale of the population dynamics.

### 3.2 Method

Genkai-Kato and Yamamura [26] examined the basal model without a PS game in the predator, and showed that the stability of the system is crucially influenced by the profitability of the less-profitable prey (i.e., *ε*_2_/*h*_2_). Especially, when *ε*_2_/*h*_2_ is either very small (’inedible’) or large (’palatable’), the system is highly unstable and the paradox of enrichment is prominent. To compare my model with their result, I investigated the non-equilibrium dynamics of my system (Eq. 2). Since the trends in the stability indices were identical for all species, I calculated stability for a single species (*X*_1_). I focused on the magnitude of population oscillation as an index of extinction risk, because the population becomes vulnerable to stochastic perturbations when its density is at the bottom of the oscillation.

### 3.3 Results

Figure 2 shows the magnitude of oscillation against the profitability of the less-profitable prey *ε*_2_/*h*_2_. For simplicity, here I set *f* = *f*_1_ = *f*_2_ and *F* = *F*_1_ = *F*_2_. When the finder’s advantage *f/F* is small (i.e., *f/F* = 0.21 or 0.51), the system is always stable regardless of *ε*_2_/*h*_2_. When the finder’s advantage is large (i.e., *f/F* = 0.81), however, the system oscillates under a range of conditions as shown in the basal model without PS game [26].

**Figure 2.**
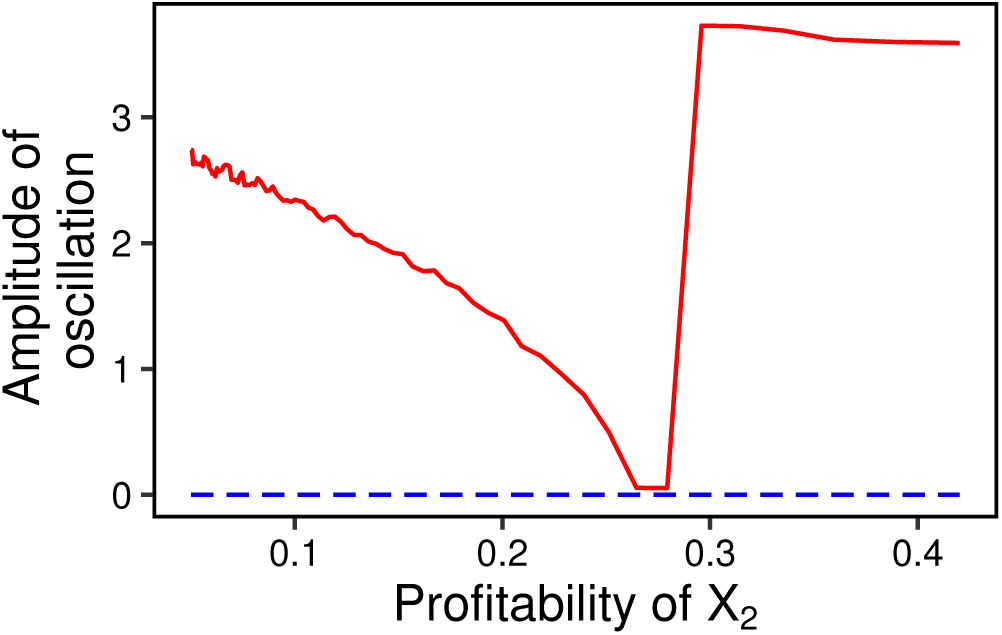
Relation between the profitability of the less-profitable prey *ε*_2_/*h*_2_ and the amplitude of oscillation, defined by the difference between the maximum and the minimum abundance of the more-profitable prey *X*_1_. Dashed line shows the cases in which the finder’s advantage *f/F* is either 0.21 or 0.51. The solid line shows the case in which the finder’s advantage *f/F* is 0.81. Numerical solution is obtained using the following parameter values: *r*_1_ = 15, *r*_2_ = 10, *a*_1_ = *a*_2_ = 1, *ε*_1_ = *ε*_2_ = 0.5, *h*_1_ = 1, *α*_12_ = 0.1, *α*_21_ = 0.4, *b* = 1, *d* = 0.25, *K*_1_ = *K*_2_ = 4, *F* = 1, *g* = 3.

Next, I considered the differences between the finder’s advantage of two prey species. I examined the magnitude of oscillation against possible combinations of *f*_1_/*F*_1_ and *f*_2_/*F*_2_, for different profitability of the less-profitable prey *ε*_2_/*h*_2_. Figure 3 shows that the system is stable under a broad range of combinations of the finder’s advantages. When *ε*_2_/*h*_2_ is small (Figure 3A and 3B), the stability of the system relies only on the finder’s advantage for the more-profitable prey *f*_1_/*F*_1_. This is because the less-profitable prey is hardly ever included in the diet because of the low profitability, and thereby *f*_2_/*F*_2_ never affects the predators’ behaviour. On the other hand, when the profitability of the less-profitable prey is large enough to actually be included in the predator’s diet, both *f*_1_/*F*_1_ and *f*_2_/*F*_2_ affect the stability (Figure 3C and 3D).

**Figure 3.**
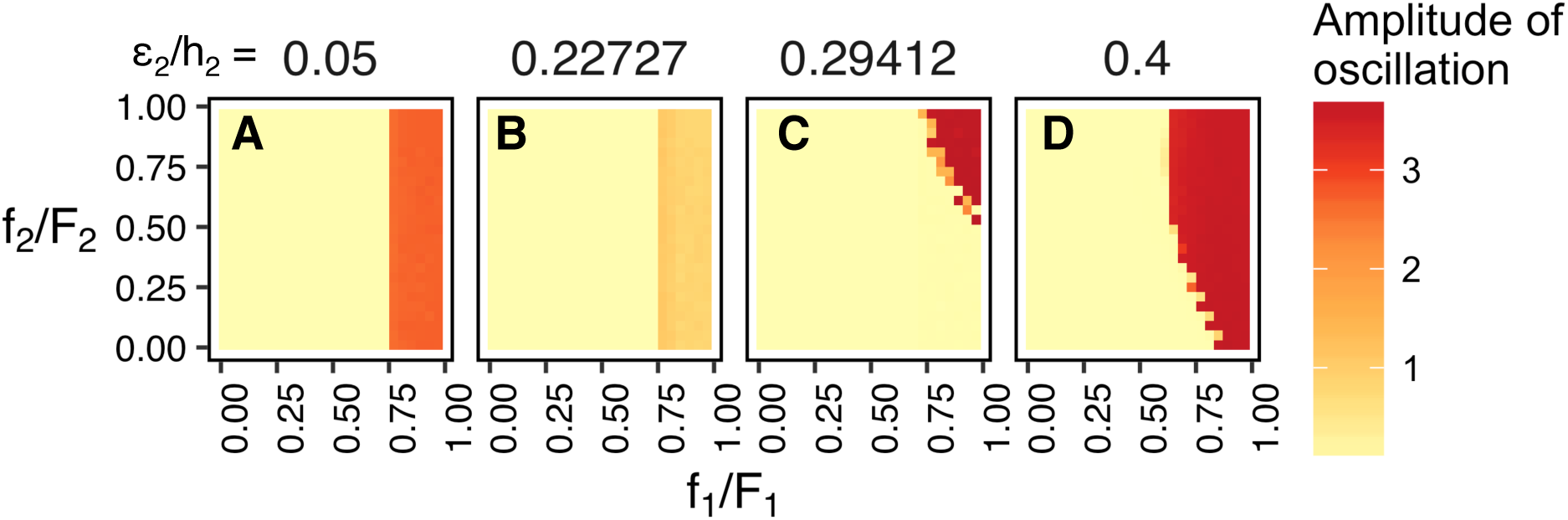
The amplitude of oscillation against the combinations of the finder’s advantages *f*_1_/*F*_1_ and *f*_2_/*F*_2_, at different profitability *ε*_2_/*h*_2_ levels: (A) 0.050 (*h*2 = 10), (B) 0.227 (*h*2 = 2.2), (C) 0.294 (*h*_2_ = 1.7), and (D) 0.400 (*h*_2_ = 1.25). The oscillation amplitude is shown in different colours from light (yellow) to dark (red). Both *F*_1_ and *F*_2_ were set to 1, and any possible combinations between *f*_1_ and *f*_2_ were tested at step size 0.05. The other parameters are the same as in Figure 2.

Finally, I investigated how the system responds to prey enrichment. For simplicity, I set *K* = *K*_1_ = *K*_2_. Figure 4A shows that the system keeps stable when the finder’s advantage is small enough (i.e., *f/F* = 0.3 or 0.5). When the finder’s advantage is large (i.e., *f/F* = 0.6), however, the system becomes unstable as *K* increases. When the system is stable, minimum densities of all three species increase with the increase of *K* (Figure 4B-D). Therefore, the paradox of enrichment is resolved when the finder’s advantage is small. On the other hand, when the system oscillates, the minimum density of the more-profitable prey *X*_1_ becomes close to zero at which stochastic perturbations would lead them to extinction. Therefore, the paradox of enrichment still exist when the finder’s advantage is large.

**Figure 4.**
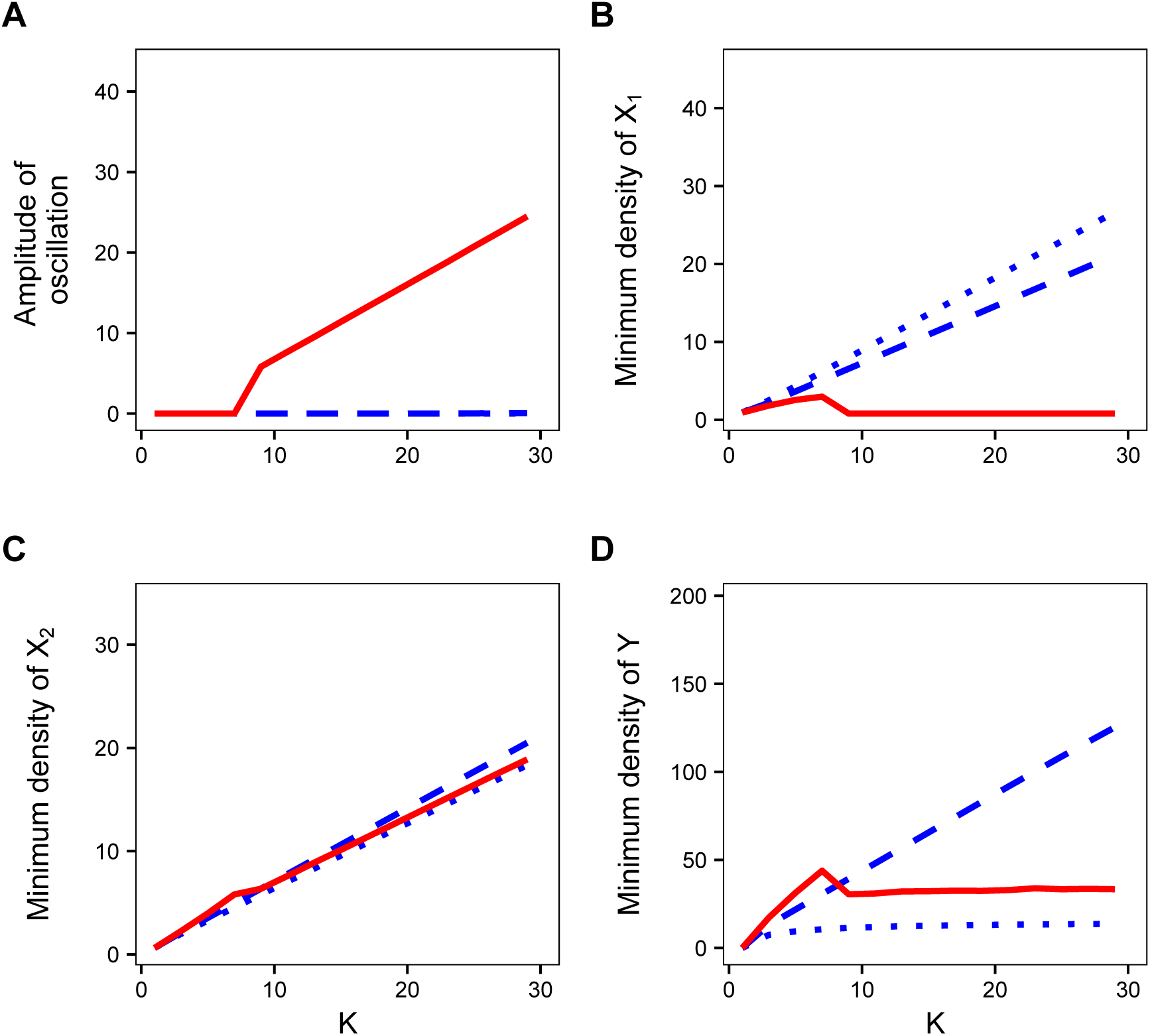
Effect of enrichment with different finder’s advantages (dotted lines: *f/F* = 0.3, dashed lines: *f /F* = 0.5, and solid lines: *f/F* = 0.6). The degree of enrichment is represented by the magnitude of the prey carrying capacity *K* (= *K*_1_ = *K*_2_). (A) Relation between enrichment and the amplitude of the oscillation. Note that the dotted line is hidden behind the dashed line. (B-D) Relation of prey enrichment with (B) the minimum density of the more-profitable prey *X*_1_, with (C) that of the less-profitable prey *X*_2_, and with (D) that of predator *Y*. The same parameter values are used as in Figure 2 except for *h*_2_, as *h*_2_ = 2.2 (i.e., *ε*_2_/*h*_2_ = 0.227).

## 4 Discussion

In this study, I demonstrated that social foraging can stabilise both a simple one-prey-one-predator system [1] and a two-prey-one-predator system [26], and thereby resolves the paradox of enrichment when scrounging behaviour is prevalent in predators. Previous studies have shown that group-living can stabilise an ecological community. For example, group formations in predators may dramatically change the functional responses, and hence stabilise predator-prey systems [35]; and resource monopolizability by higher-ranked individuals (i.e., social dominance) affects per capita food intake rates, which potentially affect population growth [36]. However, few studies have directly addressed a relation between producer-scrounger foraging dynamics and the paradox of enrichment, while scrounging may be more common than group formation or social hierarchy in animals.

For the classic Lotka-Vortella predator-prey model, Coolen et al. [24] showed that the system becomes globally stable under the existence of scrounging by predators, with-out regard to the scroungers’ proportion. On the other hand, my results show that the paradox of enrichment can be resolved only if scrounging is prevalent (i.e., the finder’s advantage *f/F* needs to be sufficiently small). This is not surprising because the stabilising effect of scrounging should overcome the destabilising effect of prey enrichment.

As Collen et al. [24] have already discussed, the stabilising effect of scrounging by predators is not new. Scrounging is one of the specific mechanisms of predator interference [11]. Predator interference, which refers to any phenomena that the per capita predator consumption rate decreases as predator density increases, is known to stabilise predator-prey dynamics (e.g. [13,14]). An intuitive explanation for this stabilising effect is that a decrease in the individual consumption rate with increasing predator density can prevent the prey population from being overexploited and hence the population oscillation can be mitigated. Similarly, the core mechanism of the stabilising effect of PS game in predators is that the proportion of producers, which actually contributes to the per capita prey capture rate, declines as predator density increases, and hence the overexploitation of prey can be avoided.

In the one-prey-one-predator system, *f/F* should be smaller than *dh/b* so as to resolve the paradox of enrichment. In reality, this inequality may be easily satisfied. Assume that prey handling time *h* becomes longer, due to an inducible defences or predator avoidance behaviour by prey for example. Making the handling time longer may also affect the amount of the finder’s advantage, because scroungers who have started to approach the captured prey in the handling time can arrive as soon as the producer started to consume the victim. Therefore, things that lengthen the handling time may reduce the finder’s advantage, resulting in raising the chance of stabilising the system. On the contrary, however, behavioural plasticity in predators may decrease a chance of stability. For example, producing predators may become more eager to consume prey so as to compensate for a loss by kleptoparasitism, which may result in reducing the prey handling time, reducing the chance of satisfying the inequality. Therefore, whether the condition of stability *f/F ≤ dh/b* is satisfied in nature or not may depend on the system, which remains open for future empirical studies.

As for the two-prey-one-predator system, the profitability of the less-profitable prey *ε*_2_/*h*_2_ affects the amplitude of oscillation as shown in the basal model investigated by Genkai-Kato and Yamamura [26] when the finder’s advantage *f/F* is large. On the other hand, the system can be stable regardless of *ε*_2_/*h*_2_ when the finder’s advantage is small (Figure 2). Interestingly, the finder’s advantages for both prey species contribute asymmetrically to the stability (Figure 3). The finder’s advantage for the more-profitable prey *f*_1_/*F*_1_ is more influential on the stability than that of the less-profitable prey *f*_2_/*F*_2_, because of the optimal diet choice by predator: The more-profitable prey is always included as a diet, while consumption of the less-profitable prey is flexible. Therefore, just keeping *f*_1_/*F*_1_ lower is enough for the system stability. When *ε*_2_/*h*_2_ is intermediate, on the other hand, keeping either *f*_1_/*F*_1_ or *f*_2_/*F*_2_ low is enough for the stability (Figure 3C). In this case, the system can be stable even if *f*_1_/*F*_1_ is high as long as *f*_2_/*F*_2_ is sufficiently small. In summary, my result suggests that scrounging behaviour does not need to be prominent in every predator-prey interaction in the community. Instead, scrounging behaviour that exists only in a subset of the community may be enough to stabilise the food-web as a whole.

I developed the PS game under multiple food types (section 3.1.2), but it should be tested empirically. Although there have been a large number of empirical studies about PS behavioural dynamics among a foragers (e.g. [22,37]), most of them were conducted under a setting of a single type of food resource. The assumption that individuals can reach the evolutionary stable state in ecological time scale rests on abilities of leaning and behavioural plasticity [29]. It is probable that animals need more memory capacity and/or cognitive skills to learn the costs and benefits associated with producing and scrounging when there are more different types of foods. Therefore, empirical tests on whether the PS foraging dynamics under multiple food types can emerge in reality is needed.

Although scrounging or kleptoparasitism is a well-documented in animal aggregations, there are many other phenomena related to social foraging I did not considered. An increasing body of empirical results show that cooperatively sharing information within a group increases foraging efficiency in a colony of social insects (e.g. [38–41]). Also, opportunities to use inadvertent social information may increase, rather than decrease, the per capita food intake rate if the environment is so uncertain that any single individuals hardly have a very accurate information, because of the “wisdom of crowds” effect or information centre [42]. Enhancing the predation efficiency may erase the stabilising effect of scrounging. Future research will clarify the relationship between such the aspects of social foraging and stability in a food-web system, and determine which stabilising or destabilising effect of social foraging is common in nature.

My study sheds light on the importance of social organisms in community ecology. Recent studies indicate that it is important for community stability to keep an interaction diversity such as intra-/inter-specific competition and mutualism, rather than merely protecting the variety of species [17]. My results suggest that it may also be important to consider whether and (if so) how the animals socially interact with other conspecifics. Ecologists should be aware of the ubiquitousness of scrounging behaviour and its potential impacts on community dynamics.

## 5 Appendices

### 5.1 Appendix A: Graphical analysis of the stability of one-prey-one-predator model

X-coordinate of the coexistence equilibrium is

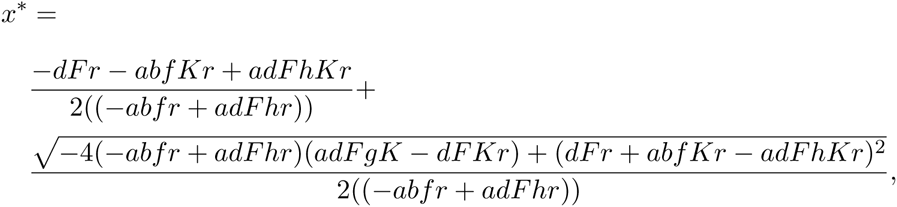

and the apex of the humped prey isocline is

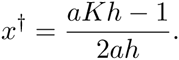

When *f/F < dh/b*, the coexistence equilibrium point is always on the right side of the hump (i.e., *x* > x*^†^), and thereby the equilibrium is locally stable. When *f/F > dh/b*, however, *x\** is larger than *x*^†^ only if prey carrying capacity is sufficiently small as *K <* (*f/F* + *dh/b*)/*ah*(*f/F − dh/b*) and when *K >* (*f/F* + *dh/b*)/*ah*(*f/F − dh/b*) the equilibrium is on the left side of the hump. Therefore, the equilibrium point becomes unstable as *K* increases. Note that whatever *f/F*, the coexistence equilibrium never exists unless *r* is sufficiently large as follows:

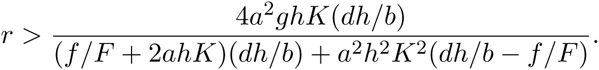

These results was obtained by using Mathematica software. The code is available from the author.

### 5.2 Appendix B: Evolutionary stability of *q\** in the PS game for multiple prey types

Consider the difference in the expected food intake *D*(*q*) = *I*_*p*_ – *I*_*s*_. By definition *D*(*q\**) = 0; and evolutionally stability requires *∂D*(*q* = *q\**)/*∂q* < 0. Differentiating and substituting, I obtain:

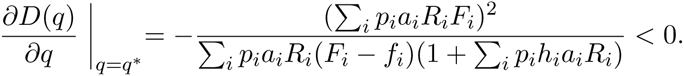

Hence *q\** is stable since (*F*_*i*_ – *f*_*i*_) > 0.

## 6 Data accessibility

Mathematica code are available upon request from the author.

## 7 Competing interest

I have no competing interest.

## Acknowledgements

Thanks to Graeme Ruxton and Kevin Laland for their valuable comments on previous versions of this manuscript.

## 9 Funding

This research was supported by Japan Society for the Promotion of Science Postdoctoral Fellowships for Research Abroad (H27-11).

## Footnotes

1 Note that in this paper I use the term “producer” to refer to a predator individual who engages in producing tactic in the PS game, instead of an energetic producer in a lower trophic level of an ecological community (e.g., plants or phytoplanktons).

## References

[1] Rosenzweig ML, MacArthur RH. Graphical representation and stability conditions of predator-prey interactions. Am Nat. 1963;97:209–223. doi:10.1086/282272.

[2] Rosenzweig ML. Paradox of enrichment: destabilization of exploitation ecosystems in ecological time. Science. 1971;171:385–387. doi:10.1126/science.171.3969.385.

[3] May RM. Will a large complex system be stable? Nature. 1972;238:413–414. doi:10.1038/238413a0.

[4] Huffaker CB, Shea KP, Herman SG. Experimental studies on predation: complex dispersion and levels of food in an acarine predator–prey interaction. Hilgardia. 1963;34:305–330. doi:10.3733/hilg.v34n09p305.

[5] Luckinbill LS. Coexistence in laboratory populations of Paramecium aurelia and its predator Didinium nasutum. Ecology. 1973;54:1320–1327. doi:10.2307/1934194.

[6] Fussmann GF, Ellner SP, Shertzner KW, Hairston NGJ. Crossing the Hopf bifurcation in a live predator-prey system. Science. 2000;290:1358–1360. doi:10.1126/science.290.5495.1358.

[7] Murdoch WW, Nisbet RM, McCauley E, deRoos AM, Gurney WSC. Plankton abundance and dynamics across nutrient levels: tests of hypotheses. Ecology. 1998;79:1339–1356. doi:10.2307/176747.

[8] McCauley E, Murdoch WW. Predator prey dynamics in environ ments rich and poor in nutrients. Nature. 1990;343:455–457. doi:10.1038/343455a0.

[9] Kirk KL. Enrichment can stabilize population dynamics: Autotoxins and density dependence. Ecology. 1998;79:2456–2462. doi:10.1890/0012-9658(1998)079[2456:ECSPDA]2.0.CO;2.

[10] Persson A, Hansson LA, Brönmark C, Lundberg P, Pettersson LB, Greenberg L, et al. Effects of enrichment on simple aquatic food webs. Am Nat. 2001;157:654–669. doi:10.1086/320620.

[11] Přibylová L, Berec L. Predator interference and stability of predator-prey dynamics. J Math Biol. 2014;71:301–323. doi:10.1007/s00285-014-0820-9.

[12] Vos M, Kooi B, DeAngelis D, Mooij W. Inducible defences and the paradox of enrichment. Oikos. 2004;105:471–480. doi:10.1111/j.0030-1299.2004.12930.x.

[13] Ruxton G. Short term refuge use and stability of predator-prey models. Theor Popul Biol. 1995;47:1–17. doi:10.1006/tpbi.1995.1001.

[14] Ruxton GD, Gurney WSC, de Roos AM. Interference and generation cycles. Theor Popul Biol. 1992;42:235–253. doi:10.1016/0040-5809(92)90014-K.

[15] Slobodkin LB. Prudent predation does not require group selection. Am Nat. 1974;108:665–678. doi:10.1086/282942.

[16] Mougi A, Nishimura K. Imperfect optimal foraging and the paradox of enrichment. Theor Ecol. 2009;2:33–39. doi:10.1007/s12080-008-0026-0.

[17] Mougi A, Kondoh M. Diversity of Interaction Types and Ecological Community Stability. Science. 2012;337:349–351. doi:10.1126/science.1220529.

[18] Giraldeau LA, Caraco T. Social foraging theory. Princeton, NJ: Princeton University Press; 2000.

[19] Barnard CJ, Sibly RM. Producers and scroungers: a general model and its application to captive flocks of house sparrows. Anim Behav. 1981;29:543–550. doi:10.1016/S0003-3472(81)80117-0.

[20] Vickery WL, Giraldeau LA, Templeton JJ, Kramer DL, Chapman CA. Producers, scroungers, and group foraging. Am Nat. 1991;137:847–963. doi:10.1086/285197.

[21] Krebs JR, Inman JA. Learning and foraging: individuals, groups, and populations. Am Nat. 1992;140:63–84. doi:10.1086/285397.

[22] Giraldeau LA, Livoreil B. Game theory and social foraging. In Game theory and animal behavior. Oxford, New York: Oxford University Press; 1998.

[23] Maynard Smith J. Evolution and the theory of games. Cambridge: Cambridge University Press; 1982.

[24] Coolen I, Giraldeau LA, Vickery W. Scrounging behavior regulates population dynamics. Oikos. 2007;116:533–539. doi:10.1111/j.2006.0030-1299.15213.x.

[25] Fryxell JM, Lundberg P. Diet choice and predator–prey dynamics. Evol Ecol. 1994;8:407–421. doi:10.1007/BF01238191.

[26] Genkai-Kato M, Yamamura N. Unpalatable prey resolves the paradox of enrichment. Proc R Soc B. 1999;266:1215–1219. doi:10.1098/rspb.1999.0765.

[27] Ma BO, Abrams PA, Brassil CE. Dynamic versus instantaneous models of diet choice. Am Nat. 2003;162:668–684. doi:10.1086/378783.

[28] Yamauchi A, Yamamura N. Effects of defense evolution and diet choice on population dynamics in a one-predator-two-prey system. Ecology. 2005;86:2513–2524. doi:10.1890/04-1524.

[29] Morand-ferron J, Giraldeau LA. Learning behaviorally stable solutions to producer–scrounger games. Behav Ecol. 2010;21:343–348. doi:10.1093/beheco/arp195.

[30] Jensen CXJ, Ginzburg LR. Paradoxes or theoretical failures? The jury is still out. Ecol Model. 2005;188:3–14. doi:10.1016/j.ecolmodel.2005.05.001.

[31] Roy S, Chattopadhyay J. The stability of ecosystems: A brief overview of the paradox of enrichment. J Biosci. 2007;32:421–428. doi:10.1007/s12038-007-0040-1.

[32] Case TJ. An illustrated guide to theoretical ecology. Oxford, New York: Oxford University Press; 2000.

[33] Charnov EL. Optimal foraging: attack strategy of a mantid. Am Nat. 1976;110:141–151. doi:10.1086/283054.

[34] Stephens DW, Krebs JR. Foraging theory. Princeton: Princeton University Press; 1986.

[35] Fryxell JM, Mosser A, Sinclair ARE, Packer C. Group formation stabilizes predator-prey dynamics. Nature. 2007;449:1041–1043. doi:10.1038/nature06177.

[36] Lee AEG, Ounsley JP, Coulson T, Rowcliffe JM, Cowlishaw G. Information use and resource competition: an integrative framework. Proc R Soc B. 2016;283:20152550. doi:10.1098/rspb.2015.2550.

[37] Giraldeau L, Soos C, Beauchamp G. A test of the producer-scrounger foraging game in captive flocks of spice finches, Loncbura punctulata. Behav Ecol Sociobiol. 1994;34:251–256. doi:10.1007/BF00183475.

[38] Sasaki T, Pratt S. Group have a larger cognitive capacity than individuals. Curr Biol. 2012;22:R827–R829. doi:10.1016/j.cub.2012.07.058.

[39] Sasaki T, Granovskiy B, Mann RP, Sumpter DJT, Pratt S. Ant colonies outperform individuals when a sensory discrimination task is difficult but not when it is easy. P Natl Acad Sci USA. 2013;110:13769–13773. doi:10.1073/pnas.1304917110.

[40] Shaffer Z, Sasaki T, Pratt S. Linear recruitment leads to allocation and flexibility in collective foraging by ants. Anim Behav. 2013;86:967–975. doi:10.1016/j.anbehav.2013.08.014.

[41] Seeley TD, Camazine S, Sneyd J. Collective decision-making in honey bees: how colonies choose among nectar sources. Behav Ecol Sociobiol. 1991;28:277–290. doi:10.1007/BF00175101.

[42] Wright J, Stone RE, Brown N. Communal roosts as structured information centres in the raven, Corvus corax. J Anim Ecol. 2003;72:1003–1014. doi:10.1046/j.1365-2656.2003.00771.x.

